# CureSCi Metadata Catalog – Making sickle cell studies findable

**DOI:** 10.1101/2021.08.13.456291

**Authors:** H. Pan, C. Ives, M. Mandal, Y. Qin, T. Hendershot, J.R. Popovic, D. Brambilla, J.K. Stratford, M. Treadwell, X. Wu, B. Kroner

## Abstract

**Objectives:** To adopt the FAIR principles (<u>F</u>indable, <u>A</u>ccessible, <u>I</u>nteroperable, <u>R</u>eusable) to enhance data sharing, the Cure Sickle Cell Initiative (CureSCi) MetaData Catalog (MDC) was developed to make Sickle Cell Disease (SCD) study datasets more Findable by curating study metadata and making them available through an open-access web portal.

**Methods:** Study metadata, including study protocol, data collection forms, and data dictionaries, describe information about study patient-level data. We curated key metadata of 16 SCD studies in a three-tiered conceptual framework of category, subcategory, and data element using ontologies and controlled vocabularies to organize the study variables. We developed the CureSCi MDC by indexing study metadata to enable effective browse and search capabilities at three levels: study, Patient-Reported Outcome (PRO) Measures, and data element levels.

**Results:** The CureSCi MDC offers several browse and search tools to discover studies by study level, PRO Measures, and data elements. The “Browse Studies,” “Browse Studies by PRO Measures,” and “Browse Studies by Data Elements” tools allow users to identify studies through pre-defined conceptual categories. “Search by Keyword” and “Search Data Element by Concept Category” can be used separately or in combination to provide more granularity to refine the search results. This resource helps investigators find information about specific data elements across studies using public browsing/search tools, before going through data request procedures to access controlled datasets. The MDC makes SCD studies more Findable through browsing/searching study information, PRO Measures, and data elements, aiding in the reuse of existing SCD data.

## INTRODUCTION

Sickle Cell Disease (SCD) is a rare hereditary blood disorder affecting an estimated 100,000 individuals in the United States [1]. The Cure Sickle Cell Initiative (CureSCi) is a collaborative, patient-focused research effort designed to accelerate promising gene therapies to cure SCD [2]. CureSCi’s Data Consortium component is creating a robust data resource for SCD researchers, with the goal of cataloguing, harmonizing, and standardizing data collection to advance science to allow for the development of safe and effective SCD curative therapies.

Using standard measures and common data elements for SCD will improve data quality and comparability. This will enhance cross-study analyses for more powerful discovery of smaller effect sizes that may influence patient outcomes or selection of effective treatments and interventions. Many initiatives and resources have been developed to promote the FAIR (<u>F</u>indable, <u>A</u>ccessible, <u>I</u>nteroperable, <u>R</u>eusable) principle to enhance data sharing and address the data silo challenge, such as the consensus measures for Phenotypes and eXposures (PhenX) Toolkit SCD Research Collection [3], the consensus recommendations for clinical trial end points developed by a partnership of the American Society of Hematology (ASH) and the US Food and Drug Administration [4,5], the series of ASH guidelines for SCD [6, 7, 8, 9], and the CureSCi Common Data Elements (CDEs) Catalog [10]. These resources provide valuable tools and resources for SCD research and trials but are not associated with study datasets.

To maximize the value of completed and ongoing studies, efforts to deposit National Heart, Lung, and Blood Institute (NHLBI)-funded study datasets of patient-level data in a centralized Data Commons at BioData Catalyst (BDC) has just started [11]. For most studies at BDC, data access is controlled through a data request process. Study metadata, including study protocols, data collection forms, and data dictionaries, provide study information about that is much more accessible than study datasets. Using metadata to describe study datasets has been recognized and promoted by the FAIR data principles [12, 13]. In this report, we describe the design and development of the CureSCi MetaData Catalog (MDC) to make SCD study datasets more Findable by curating study- and variable-level metadata and making them available through an open-access web portal. This tool does not host actual study datasets or provide the ability to query patient-level datasets. Instead, it complements the BDC so users can find information about specific data elements across SCD studies before going through data request procedures to access controlled datasets.

## MATERIALS AND METHODS

### SCD studies in the MDC

The studies included in the CureSCi MDC were initially identified as clinical and observational studies of patients with SCD that were funded by the National Institutes of Health (NIH) and, in particular, NHLBI. There was also the expectation that patient-level study data are or could be made available to the research public through a request and approval process. This initial criterion identified 14 completed or ongoing studies (Table 1). Three additional SCD registries not funded by NIH were added to the MDC because of their large sample size (i.e., the Sickle Cell Disease Association of America Get Connected Registry) or their comprehensive and longitudinal clinical and patient survey data (i.e., St. Jude’s Sickle Cell Clinical Research and Intervention Program Registry and Medical University of South Carolina’s South Carolina Sickle Cell Disease Access to Care Pilot Program). The study protocol, data dictionary, and data collection forms and basic information regarding number of patients enrolled or target enrollment, funding source, location of the data, and Principal Investigator contact information were either extracted and downloaded from BioLINCC or dbGaP, or they were requested from each study investigator and made available through the web portal.

**Table 1.**
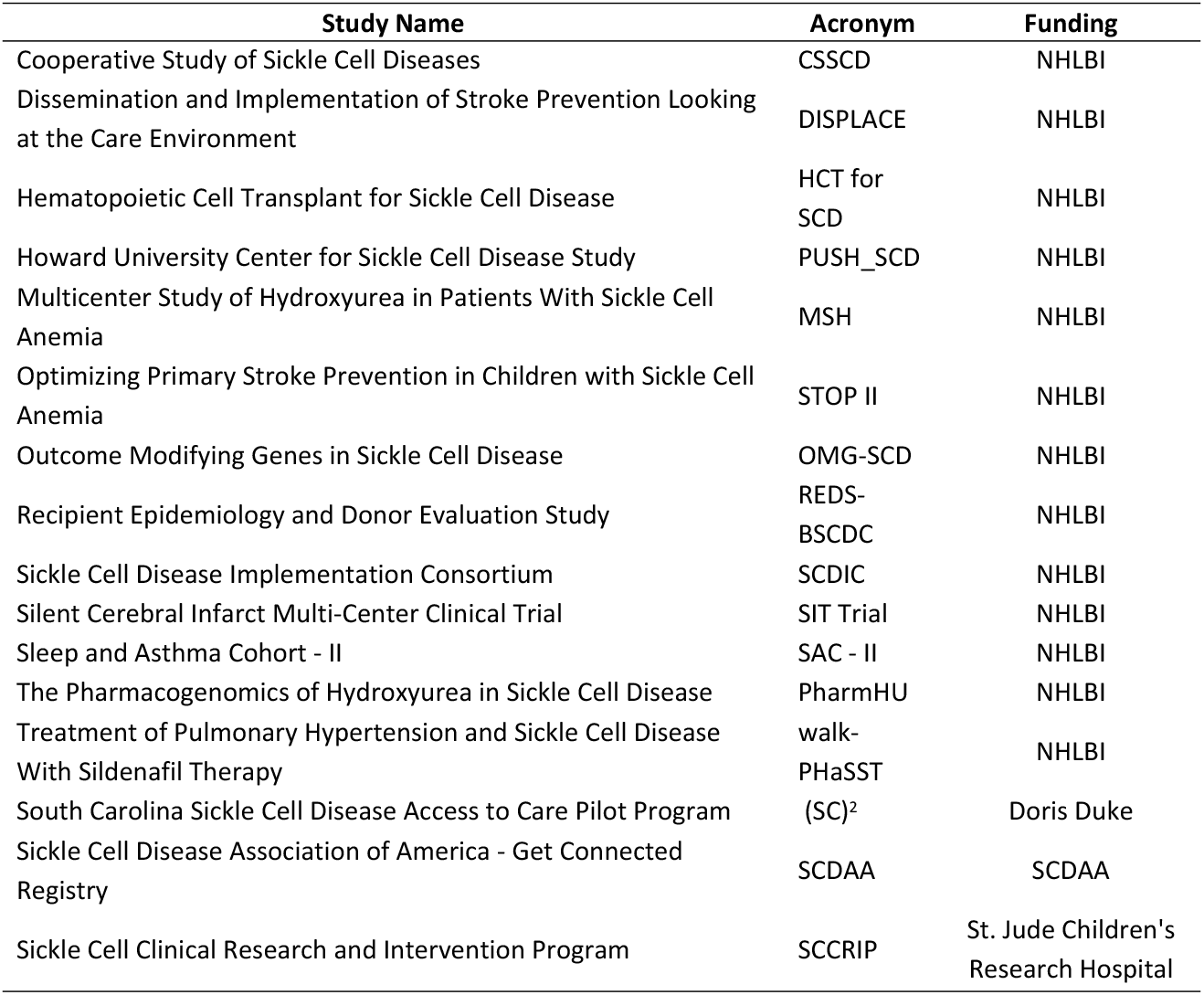
List of SCD studies released in CureSCi MDC.

### Study-level metadata curation

Key study-level metadata were curated, including 25 data fields organized into five groups as “General,” “Research,” “Access,” “Study Population,” and “Documentation.” General information, such as study design, study period, and number of subjects, was also collected. Furthermore, the studies’ focus areas and primary outcomes were summarized based on information from the literature review, the study investigator, and the ClinicalTrials.gov data resource [14]. Inclusion and exclusion criteria were obtained from consent forms, protocols, and study publications. Finally, data dictionaries, protocols, and individual data collection forms were collected, and permission to share them in the MDC was obtained.

### Data element curation

The CureSCi MDC employs a three-tiered conceptual framework to organize and curate study variables. This hierarchical classification system starts with the concept category and is followed by the subcategory and data element. Variables in each study are first cross-referenced with the existing data elements in the MDC to see if they can be categorized into existing categories. If none of the existing data elements are appropriate for the variable at hand, a new data element is created. Some study variables were excluded from the MDC, including (1) study-specific variables related to study administration, (2) variables reporting health information protected by the Health Insurance Portability and Accountability Act that could be used to identify study participants, (3) derived variables that lack documentation in the study forms preventing them from being re-derived as needed, and (4) variables not related to scientific or medical content. All variables selected by curators for exclusion were individually reviewed prior to omission from the MDC.

### Patient-Reported Outcome (PRO) Measure Curation

PRO Measures are derived from outcomes reported by patients. Prominent PRO Measurement systems that evaluate and monitor patients’ physical, mental, or social well-being that were reported in MDC-curated studies include the Adult Sickle Cell Quality of Life Measurement Information System (ASCQ-Me^®^) [15], Patient Reported Outcomes Measurement Information System (PROMIS^®^) [16, 17], Quality of Life in Neurological Disorders (Neuro-QoL™), and the NIH Toolbox^®^. These PRO Measures play a key role in assessing SCD outcomes such as pain, quality of care, and quality of life. Using the Health Measures resource [18], PRO Measures were identified and accessioned in the MDC with the measure name and source.

### Adoption of controlled vocabularies and ontologies

To increase interoperability, controlled vocabularies and ontologies such as Medical Subject Headings (MeSH) [19, 20, 21] and the Sickle Cell Disease Ontology (SCDO) [22] were referenced and adopted for initial category, subcategory, and data element creation. Domain experts were consulted regarding terms listed in the study documentation, and terms were harmonized when appropriate. Widely used study terms not included in a controlled vocabulary were included as synonyms for terms in the vocabulary. For instance, the SCDO term “SCD Related Pain” (SCDO:0001023) has nested terms for “Chronic Sickle Cell Pain” (SCDO:0000233) and “Sickle Cell Painful Event” (SCDO:0001053) that are synonymous with a pain episode.

### Developing the web-based MDC

Best practices regarding database design, data constraints, and data normalization were adopted to optimize performance, scalability, efficiency, and integrity. The CureSCi MDC web portal was developed as a fast, scalable, and interactive multi-platform web application using open-source languages such as .Net Core, JavaScript, and Bootstrap. The web portal and associated database were designed to use HTTPS, a secure and standardized communications protocol, to ensure that metadata are retrievable via the unique identifier and to maximize data accessibility. Javascript was employed to enable instant dynamic display of selected search filters/facets results.

## RESULTS

The most recent release of the CureSCi MDC (Version 2.6, June 30, 2021) incudes curated metadata from 16 studies. The studies in the MDC are quite diverse and vary with respect to study design, including clinical trials and case-control, cross sectional, registry, and cohort studies (Fig 1). Study measure distribution is quite variable with some studies being small and specific (e.g., Hematopoietic Cell Transplant for SCD) whereas others are large and comprehensive (e.g., Cooperative Study of Sickle Cell Diseases) (Fig 1B). Sifting through each study individually would be a difficult and time-consuming task; therefore, the MDC has incorporated several tools to make these studies more Findable.

**Fig 1.**
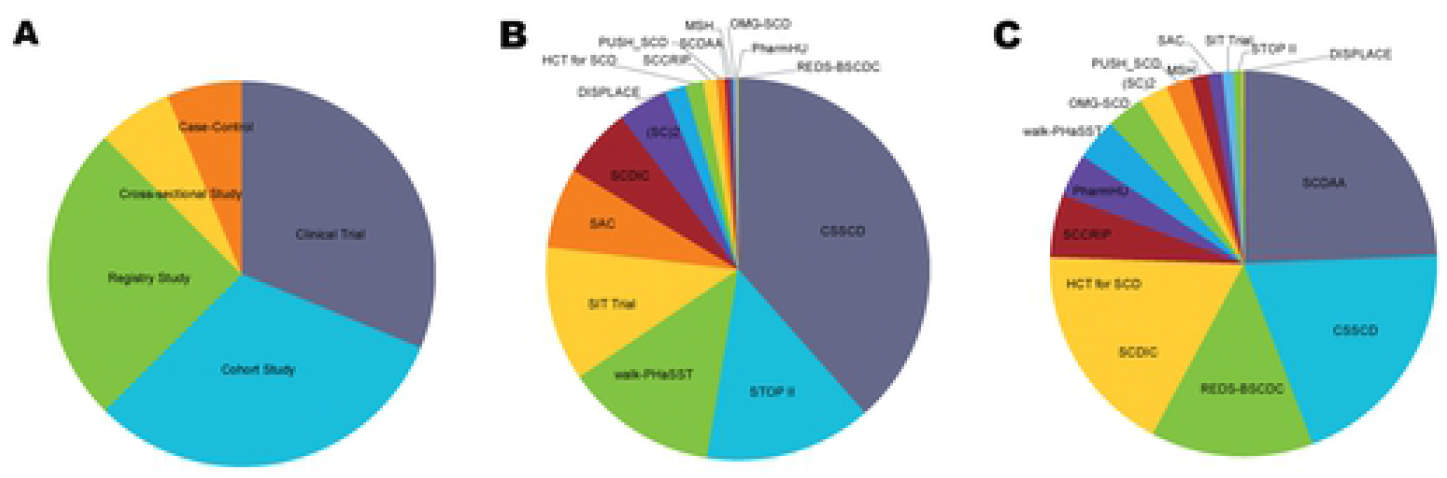
Distribution of Experimental Design, Collected Variables, and Patient Size From Studies Included in the Metadata Catalog, Available on the CureSCi MDC Homepage. A. Proportion of curated studies by study design. B. Distribution of >10,000 study variables across the 16 curated studies. C. Distribution of >20,000 subjects among the 16 curated studies.

The CureSCi MDC (https://curesicklecell.rti.org/) offers several browse and search tools. Within a study, users can browse the study-level information in 19 categories with study variables organized in the category/subcategory/data element hierarchy. Across studies, the “Browse Studies by Data Elements” tool allows users to see the number of studies that collected a given category/subcategory/data element. Two search modalities were implemented to allow queries of study-level and data element–level features, by keyword and prepopulated conceptual categories backed by the three-tiered conceptual framework. Search modalities may be used independently or in combination to enhance search capability and maximize discovery of relevant content. To achieve effective and interactive browsing, traditional search techniques were supplemented with a faceted navigation panel by grouping study variables in the three-tiered conceptual framework, allowing users to narrow down search results by applying multiple data elements filters. A tutorial of MDC features is available on the study site, https://curesicklecell.rti.org/player/CureSCi_Metadata_Catalog_v3_player.html. To date, 19 studies have been selected for curation, and data from 16 studies have already been uploaded to the web portal, as listed in Table 1.

### Browsing by study-level metadata

The “Browse Studies” tool provides access to study-level metadata for the studies included in the MDC. Detailed study information is organized into topics spanning (1) general study information, (2) research overview, focus, and outcomes, (3) access to data or biospecimens, (4) study population description, (5) individual forms and documentation available for viewing, download, or sharing via URL, and (6) data elements associated with this study. The data elements are categorized based on the three-tiered category/subcategory/data element structure. Hovering over each variable reveals a detailed description and values linked to the variable from the study forms.

### Browsing by data element

The “Browse Studies by Data Element” tool enables browsing/filtering studies that include a chosen data element selected from a user-friendly expandable menu of the category/subcategory/data element classification framework (Fig 2a). This functionality can be extended by clicking “Show Variables” to display study variables mapped to selected data elements, providing information about variable name, description, value set, and the associated study and data collection form (Fig 2b).

**Fig 2a.**
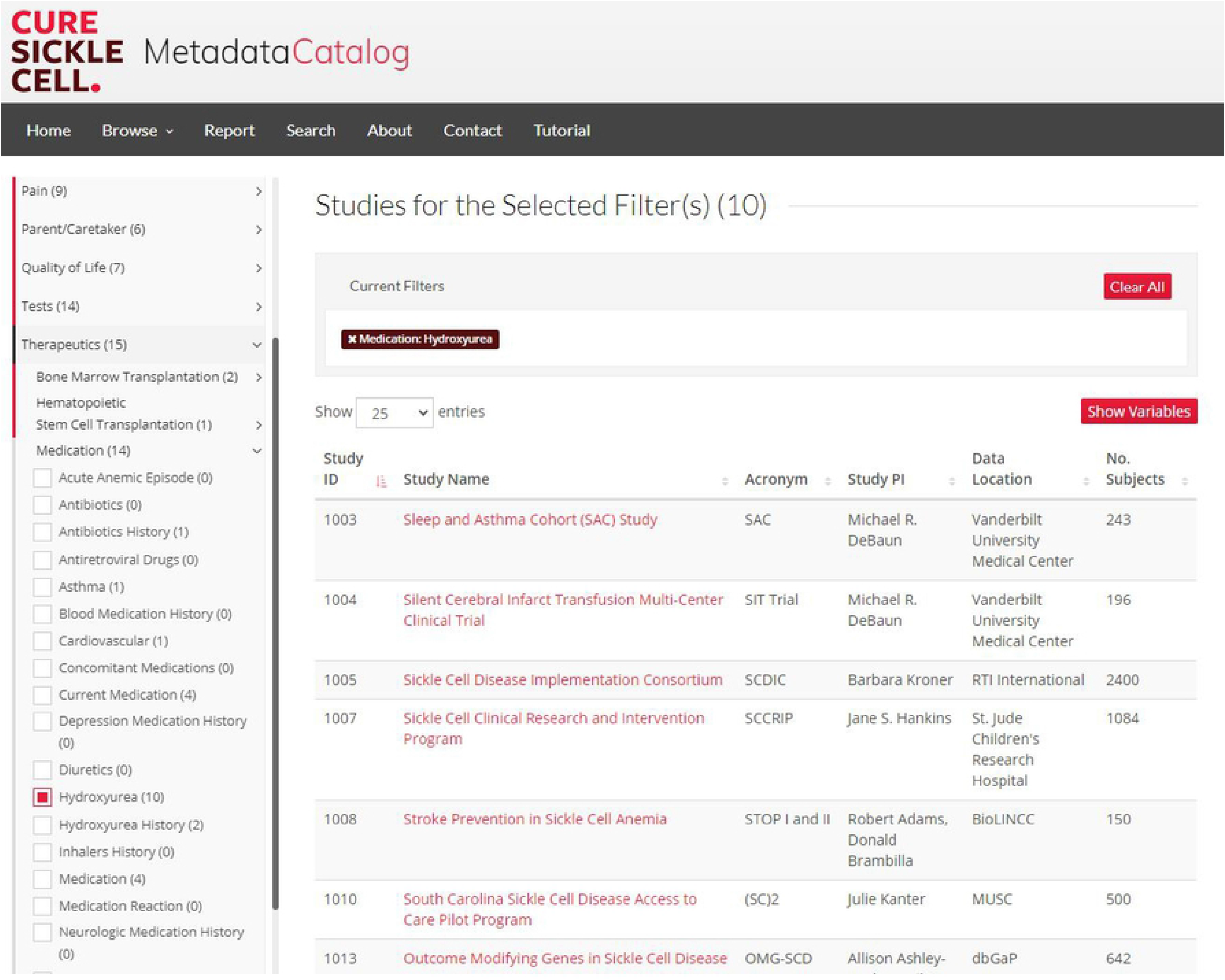
The “Browse Studies by Data Element” Tool. The tool dynamically displays studies matching the selections of data elements organized in the three-tiered classification framework.

**Fig 2b.**
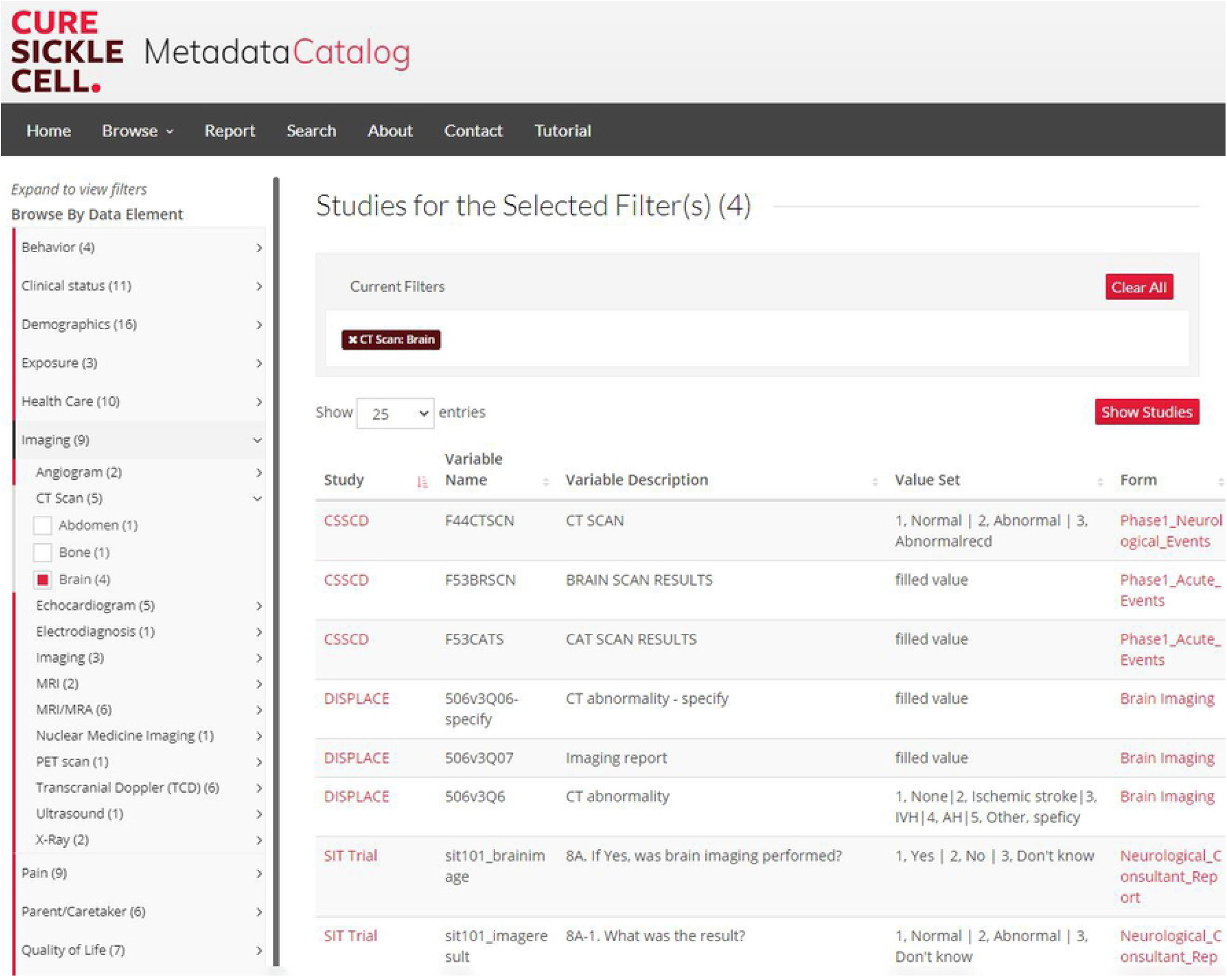
The “Browse Studies by Data Element” Tool. The “Show Variable” feature lists study variables information.

### Browsing by PRO Measure

Although data elements capture **what** is collected (e.g., depression severity), PRO Measures capture **how** the data element is collected. To date, 23 different PRO Measures across eight different PRO categories are available spanning outcomes related to functional exercise capacity, mental health, neurodevelopment, neuropsychology, pain, quality of care, and quality of life and sleep. The number of measures will increase, and categories will diversify as more studies are added to the MDC. Although indexed and searchable PRO Measure metadata fields are rarely available in public data resources, the ability to identify study datasets at the measure or instrument level reduces the potential burden of harmonization. Similar to data elements, the “Browse Studies by PRO Measure” tool enables identification of studies that measured study outcomes using selected PRO Measures (Fig 3). Furthermore, data elements and PRO Measures can be combined to refine the study search. For example, it is possible to search for studies that collected the “Pain Attack” data element AND the “ASCQ-Me Pain Episode” PRO Measure. Because many of the SCD PRO Measures were developed in recent years, PRO Measures were found only in recent studies. The power of labeling and indexing PRO Measures will be realized as additional studies adopt their use.

**Fig 3.**
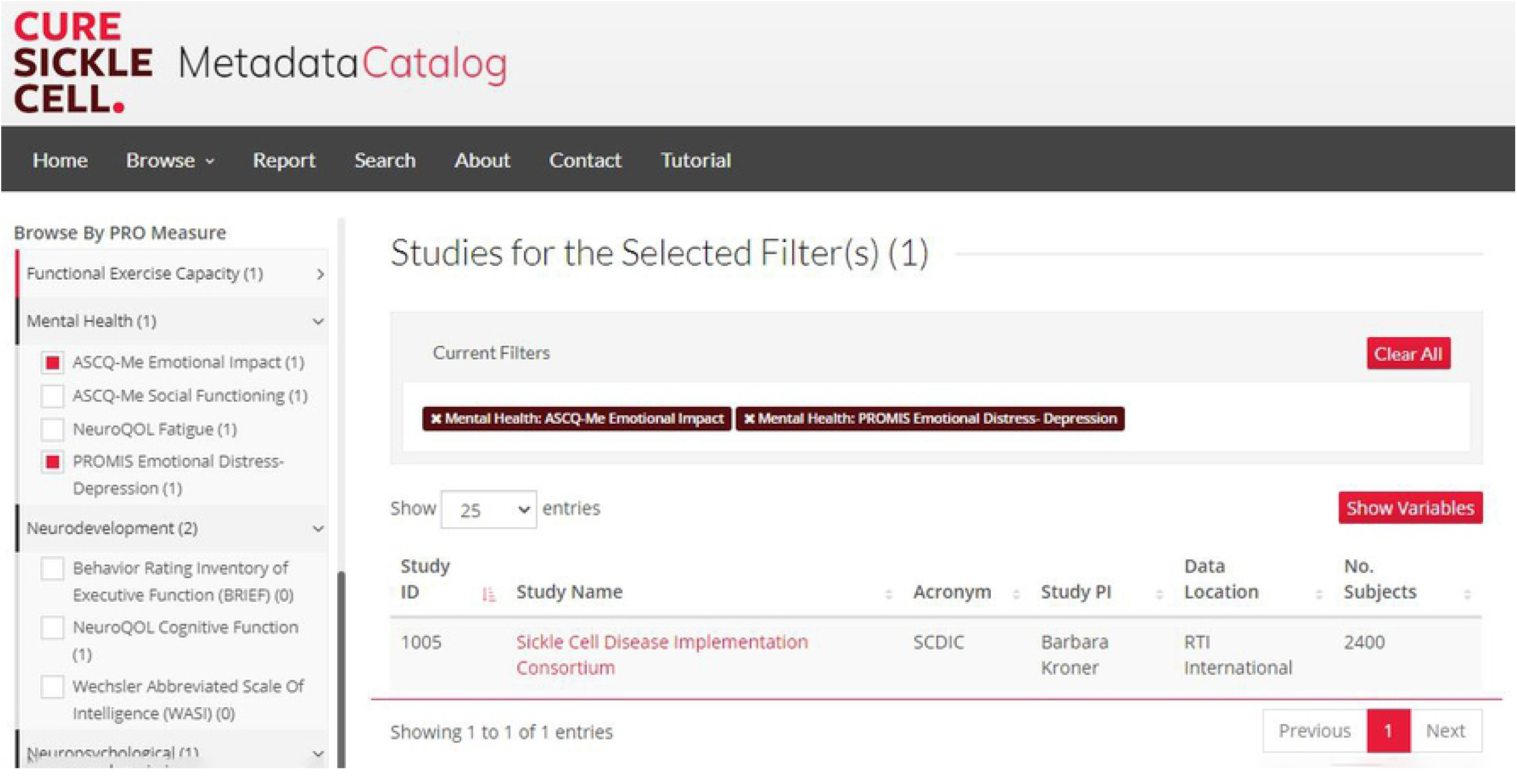
The “Browse Studies by PRO Measures” Feature Tool. This tool dynamically displays studies matching the selected PRO Measures.

### Searching with keywords or concept category filters

The CureSCi MDC includes a keyword and category search tool to help users identify relevant studies. The “Search Study Metadata by Keyword” tool allows users to search for a keyword in study name, study acronym, design, outcome, and data element, among others. The “Search Data Element by Concept Category” leverages the hierarchy of category, subcategory, or data element to select studies. For example, users interested in identifying studies collecting cardiac Magnetic Resonance Imaging (MRI) can select the “Imaging” concept category, “MRI” subcategory, and “Heart” data element. In addition, metadata keyword and data element concept category searches can be combined to refine user-defined searches further (Fig 4). One search example regards the identification of studies and data elements associated with the assessment of pain, a critical attribute in SCD studies. Terms for variables reporting pain vary with different synonyms or classification granularities, as listed in **Box 1**. Without a structured categorization, a simple keyword search for “pain” could miss data elements collected with a different synonym or granularity. To find studies that collected data for pain within the CureSCi MDC, a user can browse the “Pain” category, revealing six associated subcategories and 49 data elements for 441 variables across six studies. Searching data elements in combination with various study metadata fields can identify variables such as dactylitis that would have been missed by keyword search for “pain.”

**Fig 4.**
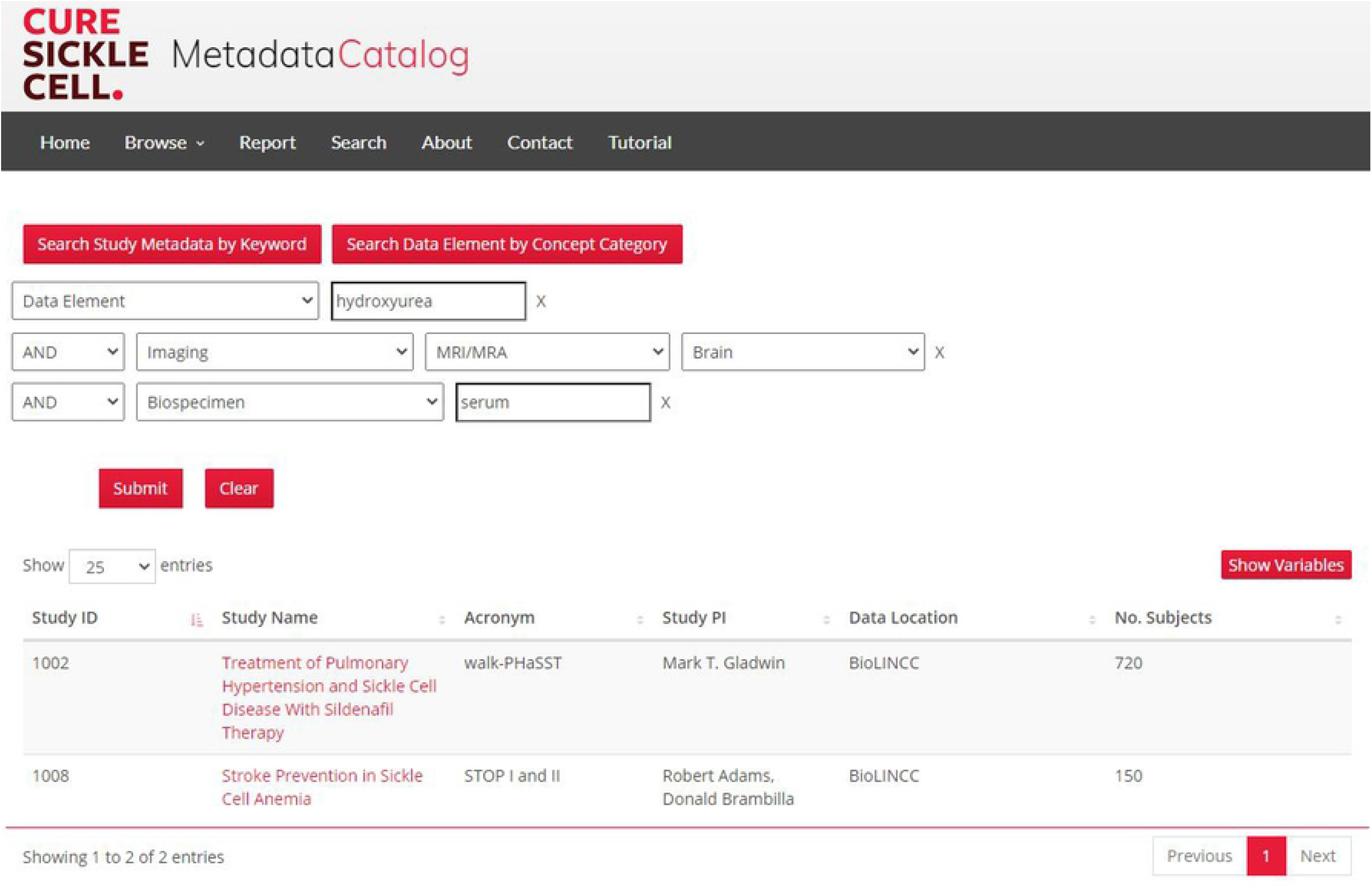
Advanced search combines options to search by keyword and prepopulated conceptual categories. Study identification can be flexibly refined by using keyword searches within a concept category/subcategory/data element or metadata field. The two search modalities may be used independently or in combination with AND/OR/NOT relations.

#### Box 1. A structured categorization of the list of terms for variables reporting pain vary with diverse names, synonyms, or classification granularities to enable browsing and search.

|  |  |
| --- | --- |
| • Abdominal Crises | • Chest Pain |
| • Abdominal Pain | • Chronic Pain |
| • Acute Anemic Episodes | • Chronic Refractory Pain |
| • Acute Chest Crisis | • Dactylitis |
| • Acute Chest Syndrome | • Headache |
| • Acute Pain | • Hyperalgesia |
| • Acute Painful Crisis | • Leg Ulceration |
| • Acute Painful Episodes | • Limb Pain and Swelling |
| • Acute Painful Events | • Neuropathic Pain |
| • Allodynia | • Pain Episode |
| • Arthritic Pain | • Painful Crisis |
| • Back Pain | • Pelvis Pain |
| • Breakthrough Pain | • Proximal Limb Bone Pain |
| • Central Sensitization Syndromes | • Respiratory Crises |
|  | • Vaso-Occlusive Crisis |
|  | • Very Severe Pain |

The multiple search features in the MDC leverage curated and indexed study and data element metadata to enable nested searches combining keywords and multiple concepts in the three-tiered concept framework. These search options provide flexibility and specificity with refined granularity. We provide a search use case to demonstrate these search capabilities, as shown in Fig 3. A search for “hydroxyurea” as a data element keyword resulted in 13 studies. Adding another search criterion of having brain MRI/Magnetic Resonance Angiography narrows the list to five studies. Adding “serum” to the search criteria narrowed the list to only two studies.

## DISCUSSION

Currently, the CureSCi MDC includes curated metadata from 16 different studies, >10,000 study variables, and data collected from >20,000 subjects. This open-access tool allows users to quickly identify collected data from historical studies relevant to current research questions, prior to going through lengthy data request procedures necessary to access the controlled datasets. Study metadata describing collected data can be browsed and searched by keyword, PRO Measures, and ontologies for single or multiple studies. Study metadata were curated to provide standard measures and common data elements through a three-tiered framework consisting of categories, subcategories, and data elements for increased findability. This framework also enables the identification of comparable study variables to support combining similar studies to increase sample size and statistical power and performing cross-study analyses. This way, the MDC provides a method to expedite research studies and data analysis resulting in improvement of the lives of people affected by SCD and supporting paths to cures.

### Unique feature of data element level curation

The primary unique feature of the CureSCi MDC is in its variable-level curation. Major NIH data repositories that host various SCD studies, including BioLINCC, BioData Catalyst, and dbGaP, provide information by curating and indexing their datasets based on study-level metadata without the ability to browse and filter at the PRO Measures or variables levels. The CureSCi MDC, a resource solely dedicated to SCD research, fills this gap by curation at the data element (variable) level. Therefore, the MDC offers the flexibility for efficient browsing of SCD research at the study, PRO Measures, and data element levels. Researchers can identify common data elements shared across multiple SCD projects and identify datasets of interest by using the nested search features offered by the MDC. It builds a bridge connecting the CDE resources and the data repositories among the studies for the SCD research community.

### Defining, classifying, and refining data elements

A significant curation challenge involves defining a set of common data elements to which study variables can be mapped and organizing the data elements in the hierarchy of categories and subcategories. Selecting a useful level of granularity for each data element is a balancing act. When granularity is too high, variables that might otherwise be meaningfully combined across studies are grouped into separate data elements, limiting the ability to pool variables from multiple studies. Therefore, we have implemented a routine systematic review to combine/expand data elements, when appropriate, to evaluate balance to allow users to find and evaluate how relevant closely related variables are for testing new hypotheses. Following consultation with experts in the SCD research community, a set of 11 concept categories and 76 subcategories were defined to which data elements were assigned. The curators perform periodic reviews for proposed changes to category and subcategory or data element placement within the hierarchy. An important action item at the conclusion of each periodic review is to update the data element assignments for previously curated studies based on the review outcomes. This ensures that search outputs are accurate because variables across all studies are mapped to the current data element definitions.

### Standardized vocabulary and ontologies

Existing standardized vocabularies and ontologies were referenced and reused during development of the three-tiered conceptual framework, primarily MeSH and SCDO. MeSH is a comprehensive controlled vocabulary developed by the National Library of Medicine for indexing and searching articles and books in the life sciences [23, 24]. It is widely used for disease and condition classification by many applications and tools such as the MEDLINE/PubMed article database and ClinicalTrials.gov. The hierarchical structure and terminology were adopted and reused in our development when applicable and becomes the backbone of our conceptual framework [23]. The SCDO is a community-driven knowledge representation system for terminology and concepts about SCD [25]. Initial efforts to adopt SCDO structure and naming were met with some challenges. Clinicians and epidemiologists reported that some terminologies, like “Abnormal Phenotype/Abnormality of Cardiovascular System, SCDO:0002245,” were awkward and inconsistent with currently used terms. Therefore, we adopted more commonly used terminologies like “Clinical Status/Cardiovascular,” and SCDO terms were included as a cross reference property for interoperability. Adaptions were also made based on the depth and significance of a concept to SCD research. For example, “Pain” is one of the most common symptoms and significantly impacts quality of life and patient ratings for quality of care. Classifying “Pain” as a subcategory under “Clinical Status” would limit the classification of data elements related to the type of pain in the established category/subcategory/data element framework. We therefore elevated “Pain” from a subcategory to the primary category level.

### Application to support data harmonization

Retrospective harmonization of existing data can be particularly challenging because it requires accessing comprehensive study documentation and data, designing a mapping approach for heterogeneous data that balances scientific precision and content equivalence with the ability to maximize data inclusion, and implementing analytic methods on harmonized data that carefully consider the potential effects of disparate study designs or populations [26]. The MDC was utilized for a pilot meta-analysis on pain assessment. MDC use enabled discovery and access to detailed study documentation and data collection forms from extant studies essential for identification of suitable studies that collected the data elements required for our analysis. The conceptual framework helped in evaluate which data elements to include based on the number of available data sets and the relationships among the data elements. This exercise demonstrated the MDC’s utility in promoting reuse of extant datasets for meta-analysis and prospective study design because it provides investigators of new studies the ability to understand how data concepts and elements were collected in past studies, thus providing tools needed to reduce and support future harmonization.

### Process enhancement

Timely curation of new studies into the CureSCi MDC will hasten and encourage data reuse such that the full potential of a research study may be realized. However, mapping study variables to common data elements is currently a manual, multi-step process requiring an intimate knowledge of both the data standards and the study variables. This process can be lengthy depending on the scope and breadth of the study. Automation of key steps involving data dictionary and case report form ingest have already shortened harmonization timelines, and continued efforts will likely result in similar efficiencies.

### Integration with other resources

Development of the CureSCi MDC is based on accessible SCD studies. This foundation provides a stable platform to expand and integrate the MDC data elements catalog and PRO Measures with well-established CDE resources, including the PhenX SCD Research Collection, the consensus recommendations for clinical trial end points developed by the ASH and US Food and Drug Administration partnership, the series of ASH guidelines for SCD, and the CureSCi CDE Catalog. This integration will provide researchers with a pivotal link between common SCD data elements with research study datasets. To facilitate dataset exchange, the MDC is building relationships and exploring methods for database interoperability with resources such as BioLINCC and BioData Catalyst, as recently highlighted in “Recent News” at BioLINCC (July 14, 2021) [27]. The MDC provides high-quality manually curated variables, along with the ontologies, which can serve as a robust training set for developing and refining Natural Language Processing algorithms to scale up metadata curation in large data repositories.

### Promotion of Data Reuse

The majority of studies included in the MDC have already concluded. Studies currently underway often delay releasing study documentation until after study results are published, which may be several years after data collection has been completed. This creates a time lag where metadata from the most recent and ongoing studies are unavailable for inclusion in the MDC, preventing development and refinement of current CDEs and promotion of more standardized data collection across newly designed studies. With the recent call for using CDEs in multiple NIH Funding Opportunity Announcements [28–33], it is now more important than ever to have an up-to-date resource to find CDE information that can streamline study design, promote dataset interoperability, and ultimately increase integration with other relevant resources. This resource depends on the collaboration and cooperation of study investigators to release their study metadata in a timely manner. Use of the CureSCi MDC will facilitate the SCD research and therapeutic community to identify studies by data element for collaborative research and promote data reuse for future SCD studies.

## CONCLUSION

The CureSCi MDC was designed to be a resource for curation of data elements and study documentation for past and current studies of SCD populations. The web-based portal presented in this manuscript provides a platform to make SCD research more Findable and promote FAIR data principles, allow investigators to browse and search metadata from existing studies to determine what data elements were collected and are common in SCD research, and finally, to identify compatible studies for cross-study analyses. The CureSCi MDC is a living entity and not a static resource. Enhancement and refinement of features will continue in response to the needs of the SCD research community and user feedback. As new study results become available, it is critical for the metadata from these studies to be incorporated into the CureSCi MDC. Providing timely access to new study documentation from the SCD community will help sustain the utility of this resource. Ultimately, the CureSCi MDC will aid in the appropriate reuse of existing data to answer pertinent research questions and the expedited design of new studies with measures and outcomes that are compatible with cross-study analysis efforts.

## Acknowledgments

The authors acknowledge the contributions of the CureSCi Data Consortium members as well as Michael DeBaun at Vanderbilt University Medical Center and Josh Richardson and Jorgen Waldermo at RTI international. The content is solely the responsibility of the authors and does not necessarily represent the official views of the National Institutes of Health.

## Authorship Contributions

All authors made contributions to the conception of the work. Qin and Pan designed the architecture; Pan, Ives, Mandal, Qin, Stratford, and Wu performed data element curation; Qin developed the web-based tools and features; Qin and Ives performed database loading; Brambilla, Hendershot, Kroner, Popovic, and Treadwell provided domain expertise on SCD and SCDIC and performed critical review.

